# Single cell micro-pillar-based characterization of endothelial and fibroblast cell mechanics

**DOI:** 10.1101/2021.10.11.463878

**Authors:** Julia Eckert, Yasmine Abouleila, Thomas Schmidt, Alireza Mashaghi

## Abstract

Mechanotransduction, the ability of cells to sense and respond to the mechanical cues from their microenvironment, plays an important role in numerous cellular processes, ranging from cell migration to differentiation. Several techniques have been developed to investigate the underlying mechanisms of mechanotransduction, in particular, force measurement-based techniques. However, we still lack basic single cell quantitative comparison on the mechanical properties of commonly used cell types, such as endothelial and fibroblast cells. Such information is critical to provide a precedent for studying complex tissues and organs that consist of various cell types. In this short communication, we report on the mechanical characterization of the commonly used endothelial and fibroblast cells at the single cell level. Using a micropillar-based assay, we measured the traction force profiles of these cells. Our study showcases differences between the two cell types in their traction force distribution and morphology. The results reported can be used as a reference and to lay the groundwork for future analysis of numerous disease models involving these cells.

## 1. Introduction

Mechanics is a fundamental property of biological cells with implications for various biological functions, ranging from single cell migration to organ-level functions, such as tissue barrier integrity regulation. The emergence of bio-printed and organ on chip models was a response to the need for modeling of mechanical alterations in numerous diseases (Evers et al., 2021). In particular, mechanical dysregulation of endothelial cells is involved in several functions and is attributed to various conditions including, autoimmune vasculopathies, viral hemorrhagic syndromes, allergic reactions, and cancer. Similar to endothelial cells, mechanical regulation of fibroblasts is involved in various cellular functions. Among these are, extracellular matrix (ECM) remodeling (Leslie et al., 2021), tissue regeneration (Li and Wang, 2011) and angiogenesis (Sorrell et al., 2007). Mechanical cues from the microenvironment modulate vascular endothelium permeability and ECM synthesis. Stresses from the microenvironment are then translated to adhesion forces created between cells, and cellular traction forces applied on the ECM (Arslan et al., 2021; Lemmon et al., 2005). Traction forces generated by the actomyosin machinery contribute to the cellular mechanical properties, and they are believed to play pivotal roles in regulating various cellular mechanosensing processes, such as cell differentiation, migration and proliferation.

Several approaches attempted to characterize the cellular traction forces. One pronounced methodology are micropillar array substrates. Through selective coating of the tips of the pillars with ECM proteins, cells are allowed to adhere and exert forces on them, which results in pillar deflections that correspond to intracellular traction forces. Here, we build upon our recent work and developed an in vitro assay to quantitatively compare the mechanical properties of two cell types, endothelial cells and fibroblasts at the single cell scale. We characterized two commonly used cell models, the human umbilical vein endothelial cells (HUVECs) and 3T3 fibroblast cells. Both are established models for studying fibroblast and vascular biology in health and disease (Medina-Leyte et al., 2020). Our results showcase discrepancies in the distribution of traction forces among the two cell types. Endothelial cells appeared to exert lower traction forces on the ECM substrate when compared to fibroblast cells. Additionally, differences in cellular morphology were observed, where a lower cell-eccentricity was detected in endothelial cells in comparison to fibroblast cell. Both cell types exert dipolar forces, however, an additional three-fold symmetry was identified for fibroblast cells in certain cell-eccentricity ranges.

## 2. Methods

### 2.1. Cell culture

3T3 fibroblast cells were cultured in high-glucose Dulbecco Modified Eagle’s Medium (D6546; Sigma-Aldrich, St. Louis, MO) supplemented with 10% fetal calf serum (Thermo Fisher Scientific, Waltham, MA), 2 mM glutamine, and 100 mg/mL penicillin/streptomycin, 37C, 5% CO_2_. For HUVECs, cells were cultured in Endothelial Cell Basal Medium 2 (PromoCell, C-22211) and supplemented with Growth Medium 2 SupplementMix (PromoCell, C-39216) and 100 mg/mL penicillin/streptomycin.

### 2.2. Immunostaining

After 22.5 h of spreading, 3T3 fibroblast and HUVEC cells were fixed for 15 min in 4% paraformaldehyde (43368; Alfa Aesar, Haverhill, MA) in phosphate-buffered saline (PBS). Furthermore, cells were permeabilized for 10 min with 0.1% Triton-X 100 and blocked for 60 min with 1% bovine serum albumin in PBS. F-actin was stained with Alexa Fluor 532-labeled phalloidin (A22282; Invitrogen, Carlsbad, CA) and the DNA with DAPI (Sigma-Aldrich).

### 2.3. Elastic micropillar arrays

Polydimethylsiloxane (PDMS, Sylgard 184) micropillar arrays of 2 μm diameter, 6.9 μm length, and 4 μm spacing in a hexagonal geometry were used for cell traction force experiments. The pillar arrays were flanked by 50 μm spacers on two sides of the array. Details of this arrangement and the experimental procedures were described earlier in detail (Hoorn et al.,2014). In brief, pillar arrays were produced on a negative silicon-wafer master made by a two-step deep reactive-ion etching process. Wafers were passivated in trichloro-silane (448931; Sigma-Aldrich). A mixture of 1:10 PDMS (cross-linker/base ratio) was poured onto the Si-master and cured for 20 h at 110C. After peeling off, the tops of the pillars were coated by micro-contact printing. For that, flat 1:30 PDMS stamps were incubated for 1 h with 40 mL of 50 mg/mL Alexa Fluor 647–labeled and 50 mg/mL unlabeled fibronectin (F1141; Sigma-Aldrich), then washed and dried. Subsequently, the stamps were gently loaded onto the ultraviolet-ozone-activated micropillar arrays for 10 min. After stamping, the arrays were passivated with 0.2% Pluronic (F-127, P2443; Sigma-Aldrich) for 1 h, and washed in PBS.

### 2.4. Microscopy

Samples were imaged at high resolution on a home-build optical microscope setup based on an inverted Axiovert200 microscope body (Carl Zeiss, Oberkochen, Germany), a spinning disk unit (CSU-X1; Yokogawa Electric, Musashino, Tokyo, Japan), and an emCCD camera (iXon 897; Andor Labs, Morrisville, NC). IQ-software (Andor Labs) was used for setup-control and data acquisition. Illumination was performed using fiber-coupling of different lasers (405 nm (CrystaLaser, Reno, NV), 514 nm (Cobolt AB, Solna, Sweden), and 642 nm (Spectra-Physics Excelsior; Spectra-Physics, Stahnsdorf, Germany)). Pillar arrays were placed upside down onto 25 mm cover glasses and inspected with an EC Plan-NEOFLUAR 40 1.3 Oil Immersion Objective (Carl Zeiss).

### 2.5. Image analysis

Images of single, nonoverlapping and randomly selected cells within the field of view of 176 x 176 mm were analyzed using MATLAB scripts (MATLAB R2018a; MathWorks, Natick, MA). Pillar deflections were quantified as previously described in detail (Hoorn et al.,2014). Deflected pillars caused by cell traction forces were distinguished from the background. The background was determined from an undeflected area of the pillar array by selecting a pillar region outside the cell area. Pillar deflections underneath the cell within the background range were discarded.

The cell spreading morphology was characterized by the moment of inertia of an ellipse using the Regionprops function in MATLAB. In respect to the minor axis of the ellipse, we measured the angular position (0°to 360°) of the deflected pillars around the nucleus. The angular position 0°was chosen in such a way that most deflected pillars were close to the major axis at 270° and less at 90° (Fig 2D).

## 3. Results

### 3.1. Endothelial cells apply less traction forces compared to fibroblast cells

Previously, we have shown that the total traction force of single 3T3 fibroblast cells exerted on fibronectin-coated micropillars is proportional to the number of deflected pillars (Eckert et al., 2021). To validate this behavior for endothelial cells, we measured the traction force of 133 HUVEC cells (Fig.1(A)). The force correlated highly to the number of deflected pillars per cell with a correlation coefficient of r = 0.9 (Fig.1(B)). The linear dependence between the number of deflected pillars and the total traction force results in a single parameter for cellular traction force characterizations, the mean traction force per deflected pillar. For HUVEC cells, we measured a mean traction force per pillar of 6.9 ± 1.9 nN. This value is significantly smaller than the averaged force per pillar applied by 3T3 fibroblasts that we have earlier reported to be in the range between 10.4 ± 3.4 nN and 15.1 ± 7.8 nN (Eckert, 2021). We should note, that the number of deflected pillars for the HUVEC cells correlated with the total number of pillars per cell (r = 0.7), i.e. the cell spreading area (Fig.1(C)), a result that we reported also for fibroblasts earlier (Eckert et al., 2021).

**Fig.1.**
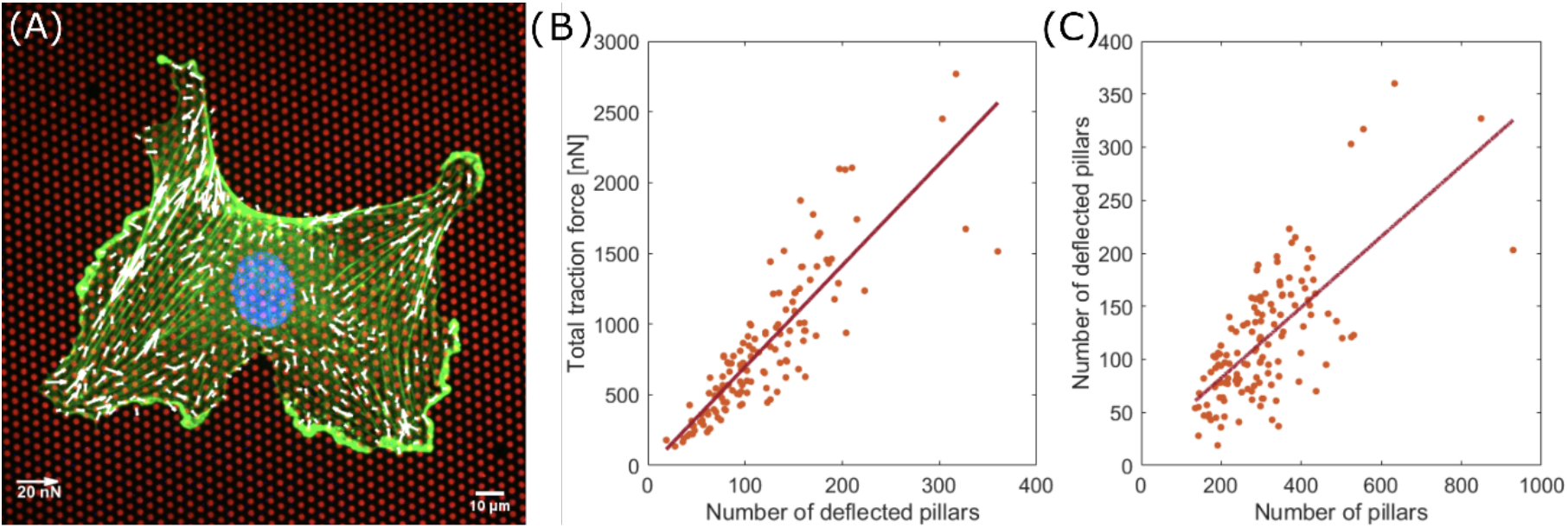
Traction force of endothelial cells increase with adhesion area. (A) Endothelial cell on fibronectin-coated micropillars (red) showing F-actin (green), the nucleus (blue) and traction forces (white). (B) Total traction force per cell as a function of the number of deflected pillars. (C) Number of deflected pillars per cell correlated with the number of pillars per cell.

### 3.2. Averaged force dipole distribution is independent of cell type

It is known that non-rounded cells generate force dipoles due to the contractility of their actomyosin machinery (Dembo et al.,1999; Schwarz et al.,2002). In round-shaped cells forces are uniformly applied on substrates and mainly distributed at the cell-periphery (Abuhattum et al., 2015). We measured the positions of deflected pillars according to the cell morphology. The cell spreading morphology was characterized by the moments-of-inertia of the cell shape approximated by an ellipse (Fig.2(A) endothelial cells, (D) fibroblasts). With respect to the minor axis of the ellipse, we measured the angular position of the deflected pillars around the nucleus (Fig.2(B) endothelial cells, 15682 deflections, (E) fibroblasts, 8824 deflections). Both distribution of the angular positions for 133 analyzed endothelial cells and 323 fibroblasts show two peaks at an angular distance of 180°, hence located at the ends of the major axes. Together with the high eccentricities of both cell lines (Fig.2(C) endothelial cells, (F) fibroblasts), the data show that both, endothelial cells and fibroblast cells, generally form force-dipoles. It should be noted that the distributions around the main peaks in endothelial cells are broader compared to those of fibroblasts, indicating a difference in the cell spreading morphology. Endothelial cells are less elongated and tend to be rounder (Fig.2(C)). In contrast, fibroblast cells are more elongated (Fig.2(F)) and form narrow force dipoles (Fig.2(E)), which can be seen from the high probability distribution for larger eccentricities.

**Fig.2.**
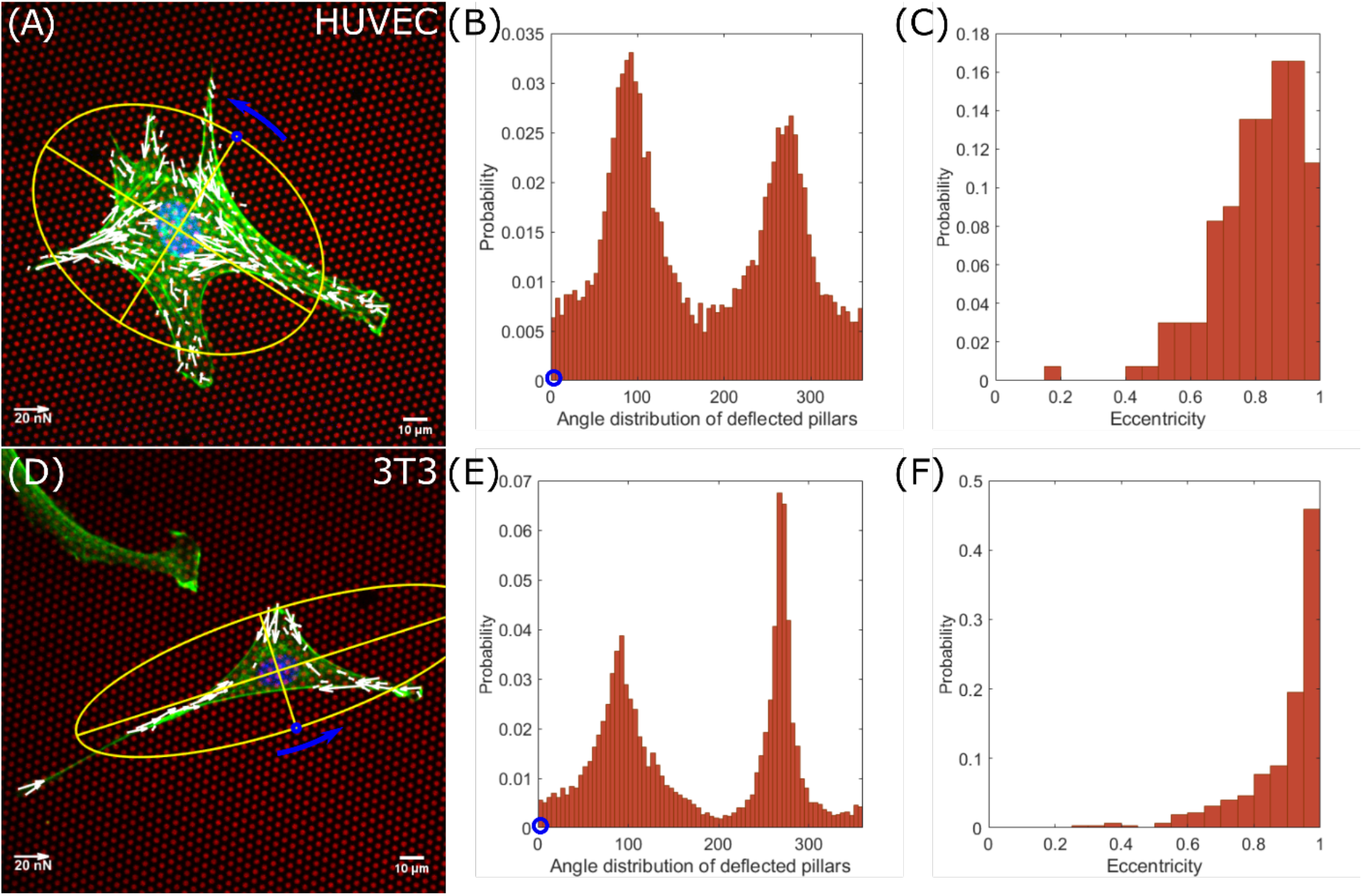
Endothelial cells and fibroblasts have dipolar force distributions. Angular distribution of deflected pillars and morphology analysis of endothelial cells (A-C) and fibroblast cells (D-F). (A,D) The cell spreading morphology was characterized by the moment of inertia of an ellipse. (B,E) Distributions of deflected pillars were assigned by counterclockwise rotation around the nucleus, starting at the short axis. (C,F) Probability distributions of eccentricities.

### 3.3. Force pole is cell morphology and cell type dependent

As a next step, we investigated the dependence of the force distribution on the cell morphology in more detail. We compared the polarity of endothelial cells and fibroblast cells with their eccentricity for similarities and differences (Fig.3(A-C) endothelial, (D-F) fibroblast). First, we investigated whether we could subtract the triangular shape of fibroblast cells from our data (Abercrombie, 1978). We plotted the angular distribution for different eccentricities, *ε*, and identified three peaks at an eccentricity range between 0.8 and 0.9 (Fig.3(E), orange arrows, 54 cells, 1650 deflections), indicating a three-pole force distribution. In comparison, very elongated cells, 0.9 < *ε* <= 1,that exerted mainly traction forces on their major axes, and that formed sharp force dipoles (Fig.3(F), 211 cells, 5691 deflections). Endothelial cells, in contrast, retained their dipole distribution in all eccentricity-ranges, even when the distribution became more uniform for rounder cells (0.8 < *ε* <= 0.9: 40 cells, 5436 deflections; 0.9 < *ε* <= 1: 37 cells, 3888 deflections.

**Fig.3.**
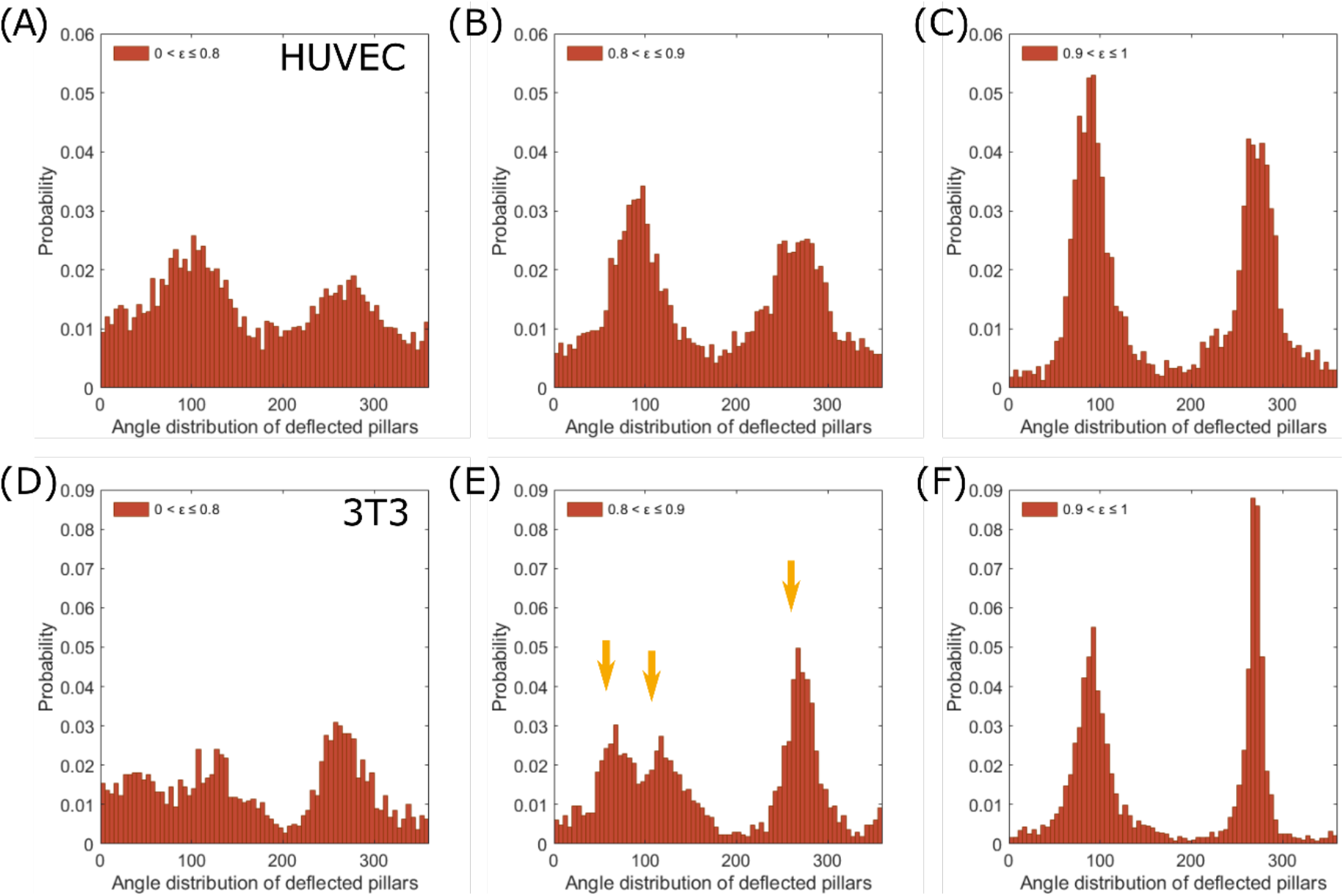
The pronounced force distribution depends on the eccentricity of cells. Angular distribution of deflected pillars according to different eccentricities, *ε*, of endothelial cells (A-C) and fibroblast cells (D-F). Endothelial cells retain two distribution peaks for all shown eccentricity ranges. Fibroblast cells show three deflections peaks at 0.8 < *ε* <= 0.9 (E) and two peak at 0.9 < *ε* <= 1 (F).

## 4. Discussion and Conclusion

Cellular mechanics is far from being homogenous across different cells. Mechanical heterogeneities of cells have been reported to largely influence several cellular processes, including response and resistance to treatment, mechanotransduction and tumor metastasis (Li et al., 2019). As such, studying the mechanical properties of individual cells is a prerequisite to provide relevant insights into the prevention and treatment of disease. Here, using a micropillar-based assay, we quantitively describe the traction forces exerted by single endothelial and compared those to fibroblast cells. We identified particular differences in cellular morphology and in the correlation of morphology with the respective force distribution. Endothelial cells we found to exert overall lower traction forces on the ECM substrate (mean traction force of 6.9 ± 1.9 nN) when compared to fibroblasts (10.4 ± 3.4 nN and 15.1 ± 7.8 nN). This may result from endothelial cells seeking cell-cell connections, necessary for network formation, and hence resulting in stronger cell-cell adhesion forces rather than cell-substrate adhesion. However, endothelial network formation relies on a balance between cell-cell and cell-substrate force interactions (Califano and Reinhart-King, 2010). While, in the case of fibroblast cells, stronger cell-substrate interactions are expected, given that fibroblasts are known to be largely responsible for ECM synthesis and remodeling (Rhee and Grinnell, 2007). We further identified a significant heterogeneity in the mean force per pillar in both endothelial and fibroblast cells. The heterogeneity in cell binding to the ECM as dictated by specific cell-ECM interactions will contribute to specific ECM remodeling, that is subsequently resulting in the structural heterogeneity of the ECM (Malandrino et al., 2018).

Regarding to the cellular morphology, individual endothelial cells appeared to be more circular in their morphology when compared to fibroblasts. These morphological differences, quantified here as the cells eccentricity, we found to be reflected in the angular force distribution as well. A broader force distribution pattern was detected in case of endothelial cells when compared to fibroblasts, as a consequence of the difference in cell spreading morphology. Both cells exhibited a dipolar force distribution, which corroborates previous studies, that migrating cells (e.g., endothelial cells and fibroblasts) demonstrate a dipolar behavior (Mandal et al., 2014). In addition, we observed a tri-polar force distribution for fibroblasts at a narrow eccentricity range (0.8 to 0.9). We speculate that the correlation of the force distribution-pattern with cellular morphology could lead to guide the directionality of cell movements, an insight that may be important for the mechanism of cell migration.

In conclusion, the micropillar-based assay reported here provides a reference for single-cell mechanical data for two commonly used cell types, HUVEC cells and 3T3 fibroblasts. We believe that our results will be of interest for future studies on the mechanics of complex tissues and organs involving these cell models. Recent studies showed a possible link between endothelial cells and fibroblasts functions in events like, inflammation and wound healing (Slany et al., 2016; Tefft et al., 2021). Thus, this assay and our data can be extended further to include cell-cell interaction to help understand how endothelial cells and fibroblasts interact and coordinate their dynamics during said events.

## Author contribution statement

JE: cell culture and sample preparation of 3T3 cells, microfabrication of micropillar arrays, data analysis. YA: cell culture and sample preparation of HUVECs. JE, YA, AM: manuscript writing. TS and AM: supervising the project. All authors: reviewing and editing the manuscript. All authors gave final approval for publication.

